# Cytoplasmic organization promotes protein diffusion

**DOI:** 10.1101/2021.07.09.451827

**Authors:** William Y. C. Huang, Xianrui Cheng, James E. Ferrell

## Abstract

The cytoplasm is highly organized. However, the extent to which this organization influences the dynamics of cytoplasmic proteins is not well understood. Here, we used *Xenopus laevis* egg extracts as a model system to study diffusion dynamics in organized versus disorganized cytoplasm. Such extracts are initially homogenized and disorganized, and will self-organize into cell-like units over the course of 20-60 min. Using fluorescence correlation spectroscopy, we observed that self-organization is accompanied by changes in protein diffusivity; as the extract organizes, proteins diffuse about twice as quickly over a length scale of a few hundred nanometers. Even though the ordered cytoplasm contained organelles and cytoskeletal elements that might be expected to interfere with diffusion, after self-organization took place, the speed of protein diffusion approached that of organelle-depleted cytosolic extracts. This finding suggests that subcellular organization optimizes protein diffusivity. The effect of organization on diffusion varies with molecular size, with the effects being largest for protein-sized molecules. These results show that cytoplasmic organization promotes the efficient diffusion of protein molecules in a densely packed environment.

## MAIN TEXT

The cytoplasm is a crowded environment filled with macromolecules and organelles (*1–3*). Its density is high enough to make intracellular motion exhibit glassy dynamics in some respects (*4, 5*). At a first glance, one might expect biochemical reactions in such an environment to be slow and inefficient. Nevertheless, protein motion in living cells is fast enough to allow reactions to occur on physiological timescales. The cytoplasm also maintains a spatial organization specific to cell identity and the cell cycle phase. One possibility is that the organization of the cytoplasm broadly affects protein diffusion and the speed of processes that depend upon diffusion, like signal transduction.

*Xenopus laevis* egg extracts are essentially undiluted cell-free cytoplasm that retains many of the biological functions of cells (*6–9*); for example, extracts can carry out cell cycles and can undergo well-organized waves of apoptosis (*10, 11*). Extracts can also be homogenized more thoroughly than intact living cells can. Moreover, recent work has shown that even well-mixed interphase extracts can spontaneously generate cell-like spatial organization from a disorganized state, forming polygonal units rich in microtubules and organelles separated by organelle-poor boundary regions (*12–14*). Here we made use of these extracts to determine how cytoplasmic organization affects protein diffusion.

We prepared an interphase-arrested cytoplasmic extract supplemented with demembranated *Xenopus* sperm nuclei and an endoplasmic reticulum (ER) dye, as described (*15*), deposited the extract in an imaging dish covered with bioinert mineral oil (*12, 16, 17*) (Fig. 1A), and monitored its organization by bright-field and fluorescence microscopy (Fig. 1B). At early time points (< 20 min), the homogenized extract showed no apparent long-range organization; organelles, including the ER, were homogeneously dispersed throughout the extract. Over the course of 20-60 min, the extract self-organized into a sheet of cell-like units (Fig. 1B). The interiors of these units were enriched in ER and other organelles; the nuclei were positioned close to the center of the interior; and separating the individual units were borders depleted of organelles. Similar results have been previously described (*12*).

**Figure 1.**
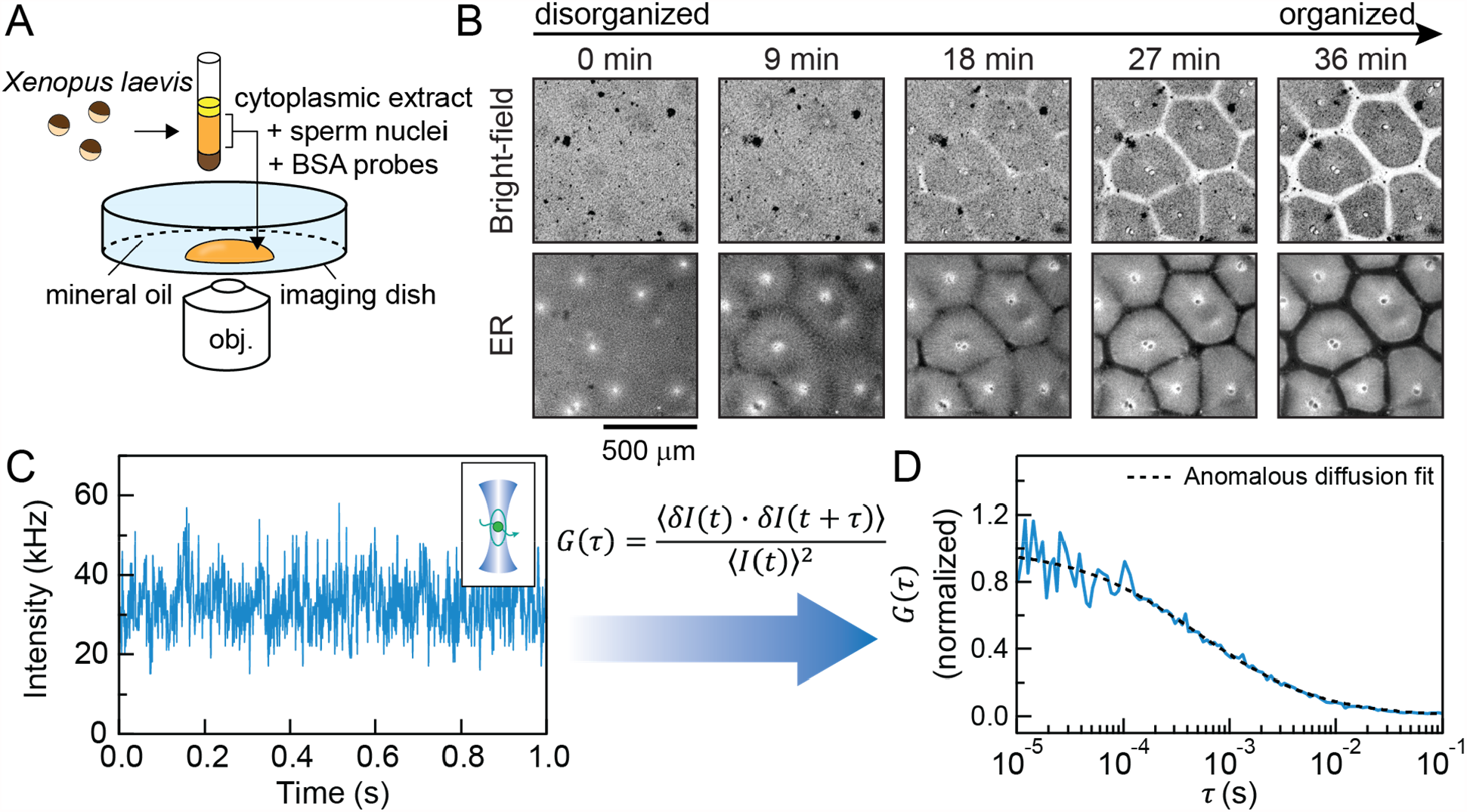
Resolving molecular diffusion in cytoplasmic extracts by FCS. (A) Schematic of the experimental setup used in this study: *Xenopus laevis* eggs were fractionated to obtain the undiluted cytoplasmic extract, which was then deposited on the surface of an imaging dish and covered with mineral oil. (B) ime-lapse images of the self-organization of cytoplasm, visualized with 1 μM ER-Red Tracker dye. (C, D) Representative FCS data for BSA-Alexa Flour 488 in organized cytoplasmic extracts (border region, as defined in Fig. 2A). The inset shows a schematic of a molecule diffusing through a confocal spot. The autocorrelation curve is fitted by an anomalous diffusion model.

We then used fluorescence correlation spectroscopy (FCS) to quantify protein dynamics in the self-organizing extract (Fig. 1C, D). FCS detects individual fluorescent molecules diffusing in and out of a diffraction-limited confocal spot; the fluorescence fluctuations enable the determination of molecular diffusion on a microsecond-to-second timescale by computing the autocorrelation function (*18*). We initially chose bovine serum albumin (BSA) labeled with Alexa Fluor 488 to represent a relatively inert protein diffusing in the cytoplasm. In a typical experiment, we first acquired confocal images to characterize the local spatial organization. We then immediately obtained FCS data to measure the local dynamics at specific locations within that field (Fig. 2A, B). Once self-organized structures emerged, we collected FCS data at locations within the border, which is depleted of ER; the outer interior, which has a contiguous ER morphology; and the inner interior, which consists of ER-depleted and ER-enriched areas (Fig. 2A).

**Figure 2.**
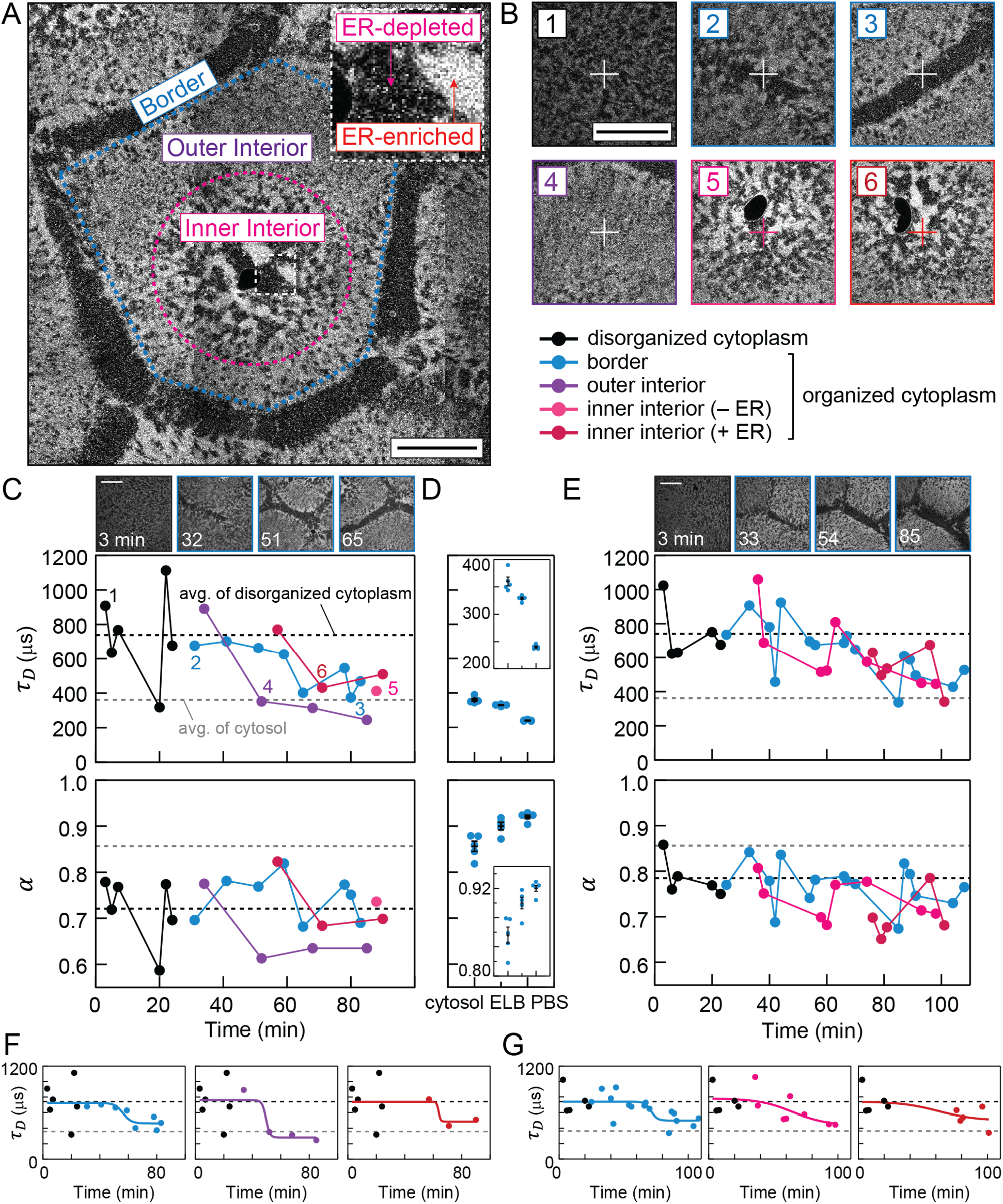
Dynamical transition in cytoplasmic extracts. (A) Spatial regions of self-organization (color-coded), classified by the ER morphology. The dark sphere near the center of the inner interior is the nuclei. The image is stitched from multiple confocal images. (B) Representative confocal images of FCS foci, which are marked by the crosshairs. The annotated numbers refer to snapshots taken prior to FCS measurements in (C). (C) Tracing protein diffusion during self-organization by FCS. The plots show the characteristic diffusion times *τ*_*D*_ and the diffusion mode *α* obtained from fitting the FCS curves. Black and grey dashed lines correspond to averages from disordered cytoplasmic extracts and cytosolic extracts, respectively. As a reference, the average diffusion times of the two conditions correspond to effective diffusion coefficients (*D*_O_) of 11.9 and 24.7 μm^2^/s, respectively. (D) Benchmark measurements from cytosolic extracts and buffers. ELB, egg lysis buffer; PBS, Phosphate-buffered saline. Insets show zoom-in of the data. Each condition is measured 5 times; error bars, SEM. (E) Biological repeat of (C). (F, G) are isolated traces from (C, E). Guide lines are the sigmoidal function, 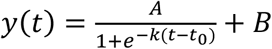, fitted to the data. Scale bars, 100 μm.

We found that an anomalous diffusion model was sufficient to describe the FCS curves measured in all cytoplasmic extracts, and we obtained the characteristic diffusion time *τ*_*D*_ and the parameter *α* (Fig. 1D, S1) (*19–21*). The value of *α* characterizes the diffusion mode as defined in the mean squared displacement (MSD) equation, *MSD* ∝ *t*^*α*^. For Brownian motion, *α* = 1, which is typically the case for BSA or dye molecules diffusing in buffers. However, molecular diffusion in complex materials like the cytoplasm is often subdiffusive, characterized by a sublinear dependence of MSD on time, with *α* < 1. This subdiffusivity has been attributed to molecular crowding and/or long-scale confinement (*1, 19–22*).

Cytoplasmic self-organization was accompanied by striking changes in protein diffusivity. Two representative experiments are shown in Fig. 2C and 2E, with a third example shown in Fig. S2. In the initially disorganized cytoplasm, the BSA dynamics were subdiffusive, and for the experiment shown in Fig. 2C, the characteristic diffusion time was 740 μs. For FCS measurements on disorganized cytoplasm from six independent experiments, the average value of *τ*_*D*_ was 677 ± 29 μs (mean ± S.E.M.). As the border regions emerged and became depleted of organelles, the diffusion time within these regions decreased by almost a factor of 2 (Fig. 2F and G). Unexpectedly, the diffusion time also decreased to a similar extent in the interior regions. Overall, the *τ*_*D*_ values for the organized cytoplasmic extracts fell to 404 ± 35 μs (mean ± S.E.M., 6 independent experiments), and there were no significant differences between the values in the border regions vs. interior regions.

We next compared the diffusion times in organized cytoplasm to those in organelle-depleted cytosolic extracts (Fig. 2D, S1; grey lines in Fig. 2C, E-G). Cytosolic extracts were prepared identically to cytoplasmic extracts, except with an additional centrifugation step that removes membrane organelles (*15*). BSA diffusion in cytosolic extracts had a characteristic time of about 360 μs with no detectable changes over a few hours. This is almost a factor of 2 faster than the diffusion time in disorganized organelle-containing cytoplasm (677 μs), and was close to the diffusion times seen in these extracts once they had become organized (404 μs). This suggests that once the cytoplasm has organized its microtubules and organelles, protein diffusion is about as fast as it can be for a solution of this protein composition and concentration. In this sense, protein diffusion appeared optimized.

Diffusion in the cytoplasmic extracts (Fig. 2C, E) was more subdiffusive than that in the cytosolic extracts (*α* ≈ 0.85; Mann–Whitney U test, two tailed p-value = 0.015; Fig. 2D, S1), suggesting that microtubules and/or organelles, even when organized, constitute barriers that hinder long-range diffusion. Overall there appeared to be a slight trade-off between *α* and *τ*_*D*_ in cytoplasmic extracts: as the cytoplasm self-organized, diffusion became faster but also slightly more subdiffusive (Fig. 3A).

**Figure 3.**
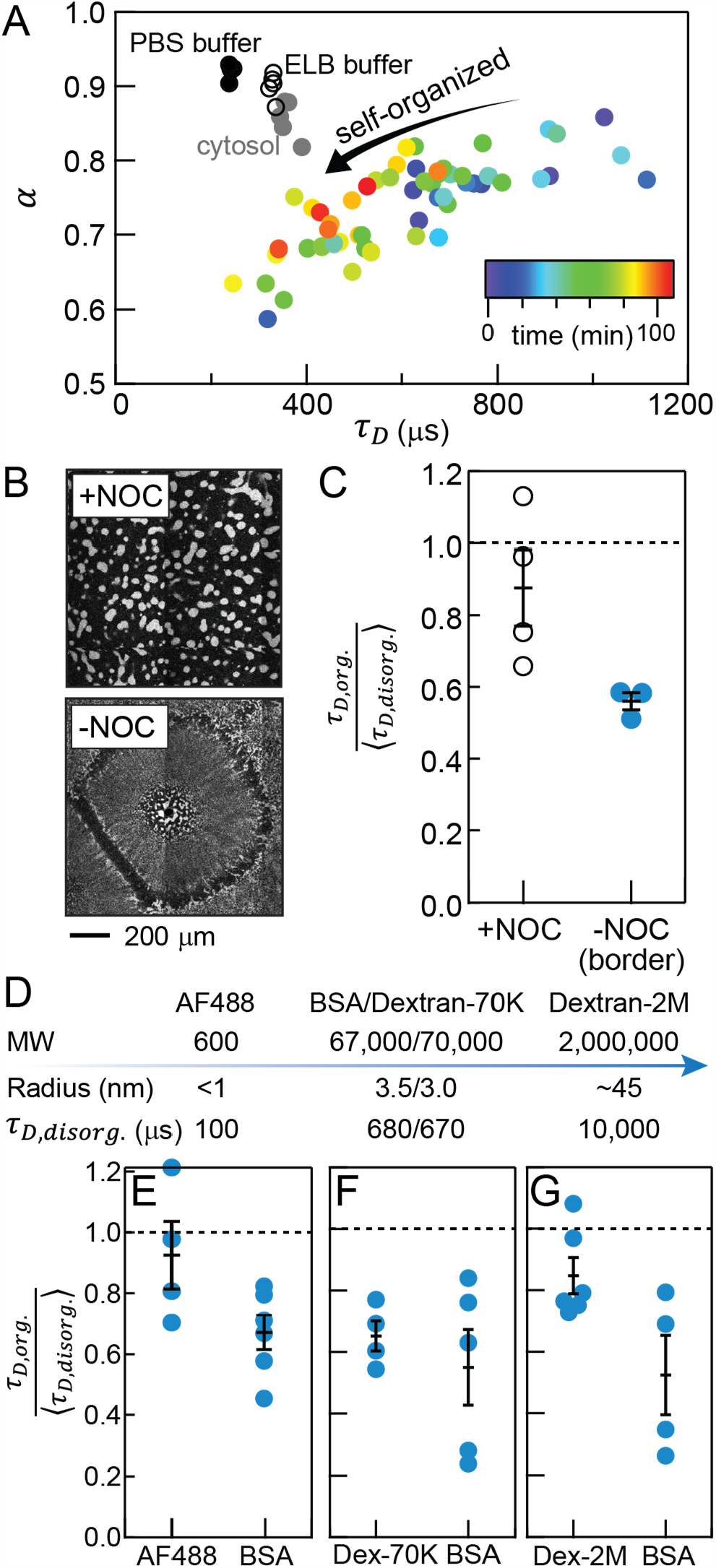
Organized cytoplasm optimizes diffusion of protein-sized molecules. (A) FCS data from Fig. 2C-E consolidated in a parameter space of {*α, τ*_*D*_}. (B) Nocodazole (NOC, 33 μM) abolishes self-organization of cytoplasm, visualized by ER-Red Tracker. (C) Nocodazole-treated cytoplasmic extracts show little changes in diffusion time (diffusion time ratio close to 1), compared with the sample without nocodazole at similar timepoints (<5 mins). The y-axis shows *τ*_*D*_ normalized by the average *τ*_*D*_ from early timepoints (<20 mins). (D) Molecular sizes of probes used in this study. (E-G) Normalized diffusion times of Alexa Fluor 488 dye (AF488), dextran-MW70,000 (dextran-70K), and dextran-MW2,000,000 (dextran-2M) in organized cytoplasmic extracts. These experiments focused on the border regions of the organized cytoplasm such that data collected <10 mins could be better compared.

Substantial variation was found in *α* and *τ*_*D*_ for different foci within the same microscopy field. This variation could be due to either error in the FCS measurements or variability in the heterogeneous extract’s dynamics. To determine how much of this variation was likely to be due to measurement error, we carried out measurements of BSA diffusion in phosphate-buffered saline (PBS), egg lysis buffer (ELB, which includes sucrose, which slows diffusion), and cytosolic extracts. The standard deviation (SD) of the *τ*_*D*_ values were about 4 (coefficient of variation CV = SD/mean = 1.6%), 5 (1.5%), and 18 (5.0%) μs, respectively (Fig. 2D, S1). Additionally, we estimated the upper bound of measurement error in cytoplasmic extracts to be ~50 μs (CV = 15%) based on repeated measurements of the same position in a mature border region (Fig. S3). The variation in our experiments (~20-30%) was larger than the measurement error (<15%), suggesting that additional variation between different positions reflects the degree of local organization. Note though that the overall trends in the data—decreases in the *τ*_*D*_ values as self-organization proceeded—were found in all three replicate experiments and in all regions of the self-organizing cytoplasm (Fig. 2C, E, S2).

As previously shown, extracts treated with the microtubule poison nocodazole do not undergo self-organization on our experimental timescale (Fig. 3B) (*12*). Correspondingly, BSA diffusion showed little change over time in nocodazole-treated extracts (Fig. 3C; statistical tests are shown in Fig. S4), indicating that microtubule function is essential for the faster diffusion.

As benchmarks, we compared the diffusion of molecules of varying sizes. We measured Alexa Fluor 488 dye and dextran-MW2,000,000 (referred to as AF488 and dextran-2M, respectively) in the disordered and ordered cytoplasm; the Stokes radii, characterized by FCS in buffers, of AF488, BSA, and dextran-2M are about < 1 nm, 3.5 nm, and 45 nm, respectively (Fig. 3D, S5) (*23*). These molecules of different sizes exhibited distinct dynamics in the cytoplasm in terms of diffusion timescales and the degree of subdiffusion. In disordered cytoplasm, the diffusion times of AF488, BSA, and dextran-2M were about 100, 680, and 10,000 μs, respectively. In terms of the diffusion mode, the intermediate-scale BSA was the most subdiffusive (*α* ≈ 0.75); dextran-2M was weakly subdiffusive (*α* ≈ 0.85) and AF488 was nearly Brownian (*α* ≈ 0.95).

Intriguingly, only protein-sized molecules were strongly affected by self-organization. Despite having greatly different timescales (100 μs versus 10,000 μs), the diffusion times of both AF488 and dextran-2M exhibited very little dependence on self-organization (Fig. 3E, G; statistical tests are shown in Fig. S4). Dextran-2M became perhaps slightly less subdiffusive in ordered cytoplasm (*α*~0.85 → 0.90; two-tailed p-value = 0.18) and AF488 remained diffusive throughout the experiments (*α*~0.95; two-tailed p-value = 0.86). To verify that molecular size was the primary determinant of the observed diffusion effects and not the molecular identity, we compared the diffusion of dextran-MW70,000 (dextran-70K) to BSA (MW 67,000) (Fig. 3F). The diffusion dynamics of dextran-70K and BSA were similar; the two probes had diffusion *τ*_*D*_ values of 670 μs and 680 μs, respectively, in disorganized extracts, and the *τ*_*D*_ values for both decreased by a factor of almost 2 after self-organization. Collectively, these results suggest that self-organization affects protein-sized molecules and has little effect on the dynamics of either small molecules or megadalton-sized molecules.

The anomalous diffusion and effective diffusion coefficients in our study agree well with live-cell measurements reported in other studies (*19, 20, 22*), suggesting that egg extracts recapitulate the physical properties of living cytoplasm. Importantly, freshly prepared extracts are homogenized, well-mixed cytoplasm, and therefore in a disordered state more thoroughly disrupted than living cells treated with cytoskeleton inhibitors. This difference may be why changes in diffusivity were not seen after inhibitor treatment in previous cell-based studies (*19, 20, 22*). It is also possible that microtubules are required to establish but not to maintain cytoplasmic structures acting as diffusion barriers (such as the ER sheets (*24*)). If so, cytoskeleton inhibitors may not fully revert these pre-established organizations.

We propose a model based on molecular and organelle reorganization as a mechanism to optimize molecular diffusivity in the ordered cytoplasm (Fig. 4A). In the disordered cytoplasm, molecules and organelles are dispersed in the cytosol such that a diffusing inert protein encounters frequent collisions; therefore, its diffusion is slow. As the cytoplasm self-organizes, molecules coalesce and form complexes with longer length-scale structures; consequently, a diffusing protein encounters fewer collisions locally, though the organized structures still hinder long-range diffusion. Effectively, organized cytoplasm creates longer obstacle-free paths, which are analogous to mean free paths in chemical kinetics, for diffusing molecules. These effects are illustrated with a mean-squared displacement diagram (Fig. 4B). Short-range diffusion in self-organized cytoplasm closely matches that in the organelle-depleted cytosol over a timescale of ~500 μs, and a distance scale of ~200 nm (corresponding to ~85% of cytosolic MSD). Conceptually, our model is related to the Lorentz model, which describes obstructed motion in a spatially heterogeneous medium (*21*). Our model describing the disorganization-to-organization effects is analogous to a higher-to-lower change in obstacle density in the Lorentz model.

**Figure 4.**
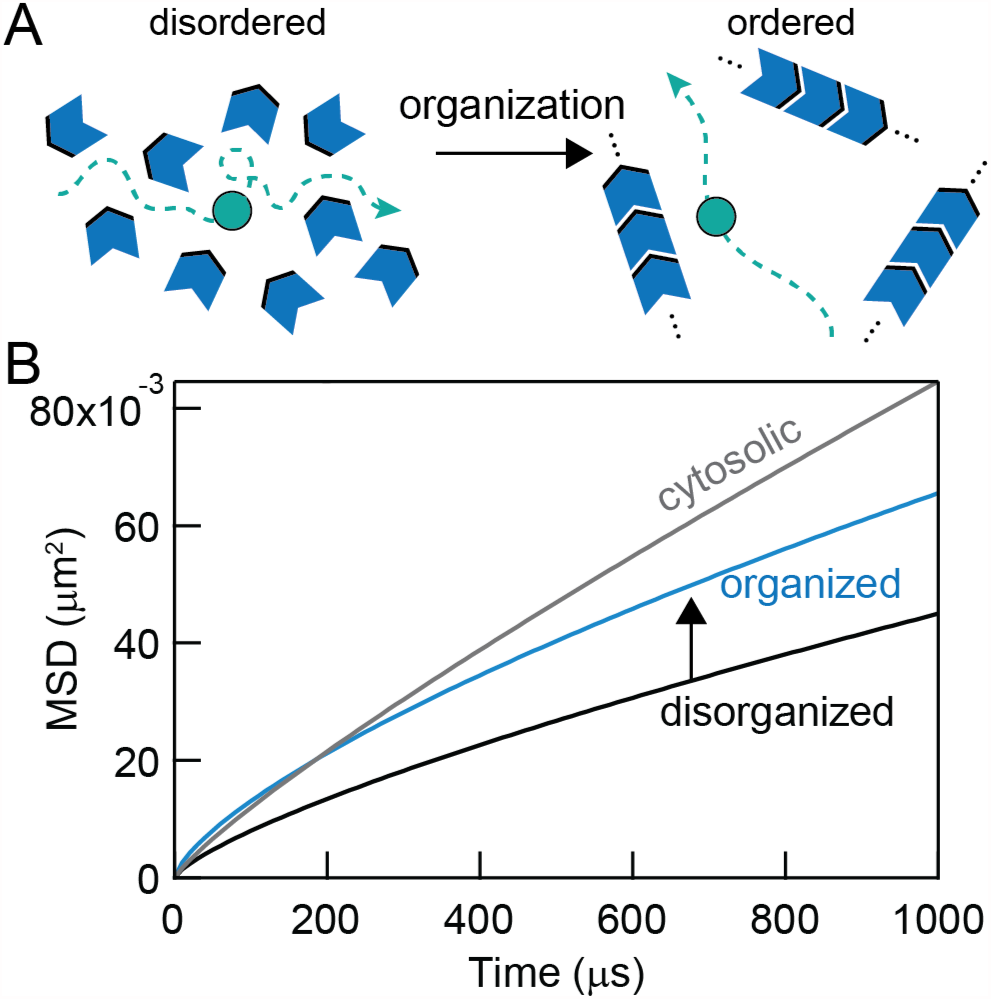
Organized cytoplasm optimizes diffusion likely via a reorganization mechanism. (A) Schematic of the model: molecular and organelle reorganization optimizes molecular diffusion by creating longer mean free paths for diffusing molecules. (B) Diffusion effects of molecular reorganization illustrated with mean squared displacement, *MSD*(*t*) = Γ*t*^*α*^, where Γ is the generalized diffusion coefficient (*21*). The parameters {Γ, *α*} estimated for disorganized cytoplasm, organized cytoplasm and the cytosol are {8, 0.75}, {8.25, 0.70}, and {30, 0.85}, respectively.

Other mechanisms may additionally regulate molecular motion in the cytoplasm, such as motor-based transport and active diffusion (*25–27*), viscosity adaptation (*28*), and phase separation (*29–31*). However, these studies, and ours, do not necessarily focus on the same length scale and timescale of cytoplasmic dynamics; our results further show that diffusion effects have size-dependence on self-organization. Nevertheless, cytoplasmic organization constitutes an additional mechanism that has substantial effects on the dynamics of protein-sized molecules.

## Acknowledgments

This work is supported by the National Institutes of Health (NIH) grant 1R35 GM131792. We thank Youngbin Lim at the Cell Sciences Imaging Facility for microscopy support, Julia Kamenz for suggestions on the experimental conditions, and Jo-Hsi Huang for experimental support. We thank Yuping Chen, Jo-Hsi Huang, and the Ferrell lab for helpful discussions and comments on the manuscript.

## Notes

### Competing Interest Statement

The authors have declared no competing interest.

