## Supplemental Information for "Cytoplasmic organization promotes protein diffusion"

### Supplementary Information for
### Cytoplasmic organization promotes protein diffusion

William Y. C. Huang[1], Xianrui Cheng[1,2*], and James E. Ferrell Jr.[1,3*]

[1] Department of Chemical and Systems Biology, Stanford University School of Medicine, Stanford, CA 94305, USA.
[2] Present Address: Department of Biological Sciences, University of Southern California, Los Angeles, CA 90089, USA.
[3] Department of Biochemistry, Stanford University School of Medicine, Stanford, CA 94305, USA.

**This PDF file includes:**

Supplementary Text
Figures S1 to S5
SI References

#### SUPPLEMENTARY INFORMATION TEXT

##### Materials and Methods

***Xenopus laevis* egg extracts.** Interphase-arrested cytoplasmic extracts and cytosolic extracts were prepared following the protocol of Deming and Kornbluth, except without the energy regeneration mix (*1-3*). Cytoplasmic extracts were placed on ice before imaging on the same day; cytosolic extracts were flash-frozen and kept in a -80°C freezer before use. Demembranated sperm nuclei were prepared as described (*2*), and added into the cytoplasmic extracts at a final concentration of about 160 nuclei/μL. ER-Tracker Red dye (ThermoFisher) was added at a final concentration of 1 μM to visualize the endoplasmic reticulum (ER). The extracts were further supplemented with cOmplete EDTA-free protease inhibitor cocktail (Roche) at the recommended concentration (1x) and incubated for 30 min on ice prior to imaging. For experiments that inhibited self-organization, we added nocodazole (MilliporeSigma) at a final concentration of 33 μM.

One of the desired diffusion probes was added to an extract right before imaging at the indicated final concentration: 25 nM albumin from bovine serum with Alexa Fluor 488 conjugate (BSA-AF488) (Invitrogen), 25 nM Alexa Fluor 488 (Invitrogen), 25 nM dextran-fluorescein-MW70,000 (dextran-70K) (Invitrogen), or 10 nM dextran-fluorescein-MW2,000,000 (dextran-2M) (Invitrogen). All probes were centrifuged at 16,000 g for 5 min to reduce aggregates before adding to extracts. Extracts were mixed thoroughly during each step. Extracts (2 μL) were then dropped on a bioinert μ-Dish 35 mm imaging chamber (No. 1.5) (ibidi) and immediately covered with heavy mineral oil (MilliporeSigma). The extracts formed similar self-organized structures in this simple setup, compared to the previous chamber setup that sandwiched the extracts with a defined spacing (*1*). We typically began imaging sessions within 3 min after extracts were taken off the ice.

***Confocal and FCS microscopy.*** All confocal images and FCS data were collected with an inverted Zeiss LSM 780 multiphoton laser scanning confocal microscope at room temperature (22°C). Samples were excited by lasers that passed through dichroic mirrors and focused by a C-APO 40x (FCS-certified) water-based objective. Signals were collected by the LSM BiG module with GaAsP photodetector, which enabled the selection of detection wavelength. For extract experiments, we typically focused the confocal spot about 30-40 μm above the dish surface, which often placed the nuclei near the focal plane. All data were acquired using the ZEN Black software.

For FCS, samples were excited by a 488 nm laser line (at 0.4% power) that passed through an MBS488 dichroic mirror. Emissions were collected by integrating signals from wavelength 499-579 nm. The pinhole was aligned using the "Adjust pinhole" function in the software. The spot-size was calibrated by averaging spot-sizes determined by fluorescein (Sigma Aldrich), Alexa Fluor 488 (Invitrogen), Atto488-carboxylic acid (Atto-Tec), and 0.1 μm TetraSpek microspheres (Invitrogen) in water at room temperature assuming known diffusion coefficients. For calibration, each FCS curve was acquired for 5 min. For extracts, each data point corresponds to an averaged FCS curve taken from six 10 s curves (a total of 1 min); in rare instances (<5%), irregular FCS curves, such as those caused by aggregates, were removed from the analysis. In the case of dextran-2M, each data point corresponds to the average of twelve 5 s curves (shorter acquisition segments enabled better removal of irregular curves distorted by aggregates while keeping sufficient signals). All averaging was calculated in the ZEN software.

For confocal images, samples were excited by a 561 nm laser line (at 2.0% power) that passed through an MBS488/561 dichroic mirror. Emissions were collected by integrating signals from wavelength 578-695 nm. The pixel time was 2.55 μs and the gain was set to 700. Each confocal image was collected by scanning 512×512 pixels, corresponding to 354×354 μm. Tile-scan combined 5×5 adjacent images, resulting in 2354×2354 pixels (1630×1630 μm).

***FCS analysis.*** The ZEN program calculated time autocorrelation functions by $G_{ZEN}(\tau) = \frac{\langle I(t)I(t+\tau)\rangle}{\langle I(t)\rangle^2}$, where $I(t)$ is the fluorescence intensity, and $\tau$ is the delay time. This function relates to our definition of autocorrelation function by $G(\tau) = \frac{\langle \delta I(t)\delta I(t+\tau)\rangle}{\langle I(t)\rangle^2} = G_{ZEN}(\tau) - 1$, where $\delta I(t) = I(t) - \langle I(t)\rangle$ (*4*). All subsequent fittings were performed in Igor Pro (version 6). FCS data were fitted with a 3D anomalous diffusion model (*5*):

$$G(\tau) = \frac{1}{N\left(1+\left(\frac{\tau}{\tau_D}\right)^{\alpha}\right)\sqrt{\left(1+\frac{1}{s^2}\left(\frac{\tau}{\tau_D}\right)^{\alpha}\right)}}$$

where $\alpha$ is the diffusion-mode parameter (as defined in the MSD equation, $MSD(t) \propto t^{\alpha}$), $\tau_D$ is the characteristic diffusion time, $N$ is the particle number, and $s$ is the structural parameter of the optics. For calibrations, the data were fitted to a Brownian model, with $\alpha = 1$. For most FCS curves, we fitted a time range of 10 μs-0.1 s; for faster (free dyes) or slower (dextran-2M) diffusion, we expanded the fitting range down to 1 μs or up to 3 s, respectively.

***Statistics and statistical tests.*** All experiments were repeated at least twice. The primary experiment measuring BSA diffusion in cytoplasmic extracts has been repeated 6 times. We used Mann-Whitney U tests (also known as the Wilcoxon rank-sum test) for all data comparison since the Mann-Whitney test is a nonparametric method that makes no assumption of the distribution. The results of the statistical tests are presented in Fig. S4. All comparisons were analyzed from experiments on the same day using the same batch of freshly prepared egg extracts.

#### SI FIGURES

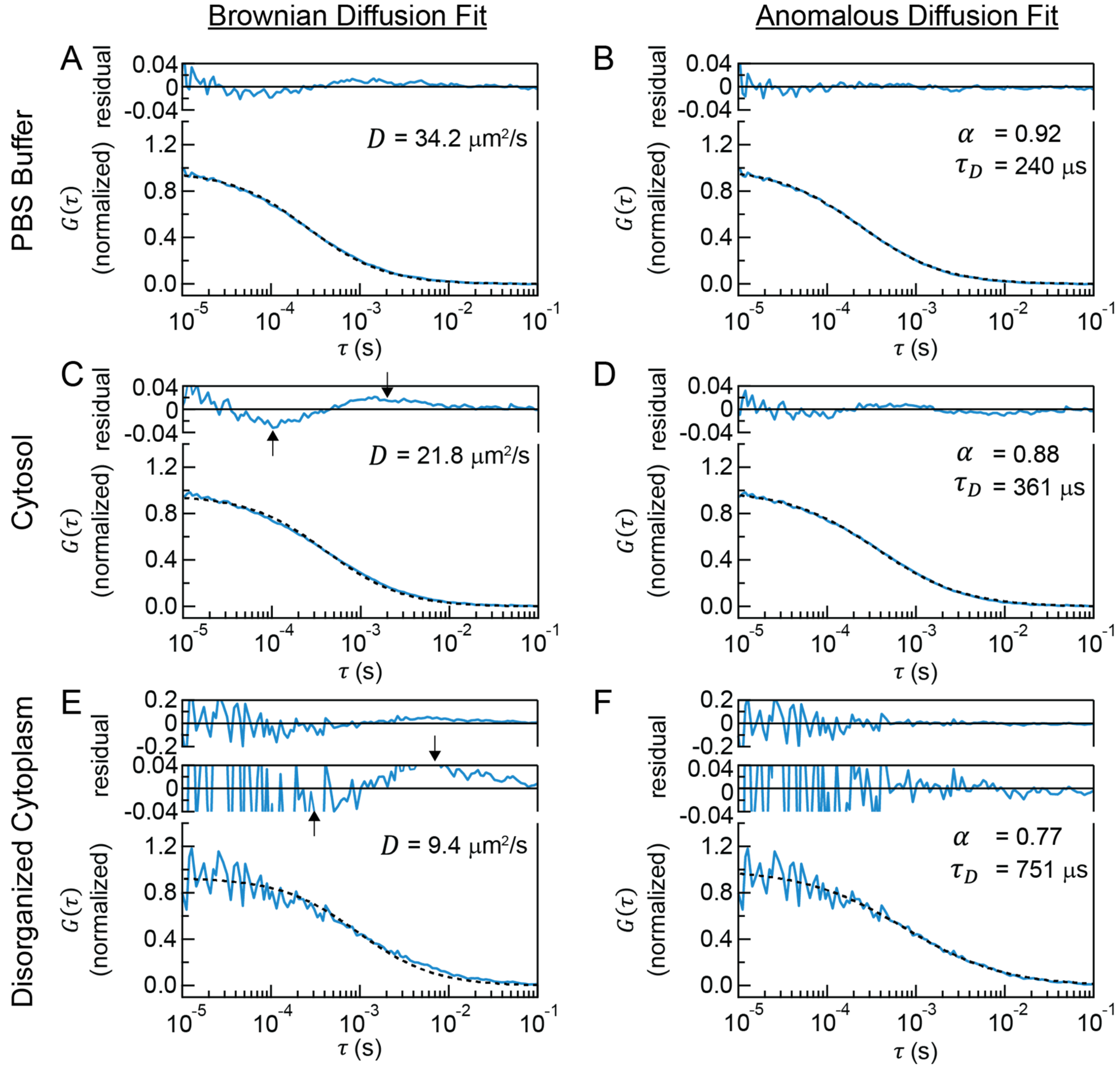


**Figure S1 Diffusion analysis of FCS autocorrelation functions**

(A, C, E) Fitting a Brownian model to data for BSA diffusion in PBS buffers, cytosolic extracts, and disordered cytoplasmic extracts. (B, D, F) Fitting an anomalous diffusion model to the same data for the same conditions. Note that Brownian motion ideally yields $\alpha = 1$ even with anomalous diffusion fits. The effective diffusion coefficients of (B, D, F) are 37.2, 24.7, and 11.9 $\mu m^2/s$, respectively.

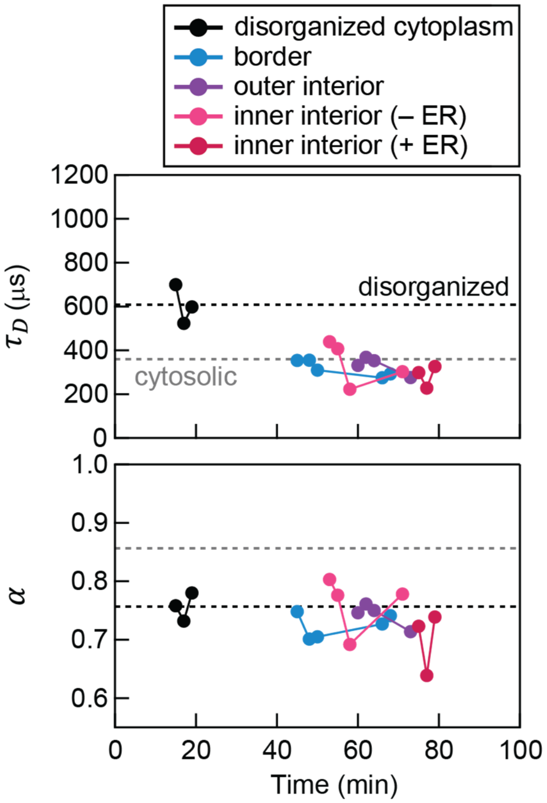


**Figure S2 Dynamical transition in cytoplasmic extracts: additional biological replicate**
A repeat of Fig. 2 experiments. In this sample, the cytoplasm formed organized structures earlier than the other experiments; as a result, the diffusion changes between the disorganized and organized states were especially apparent.

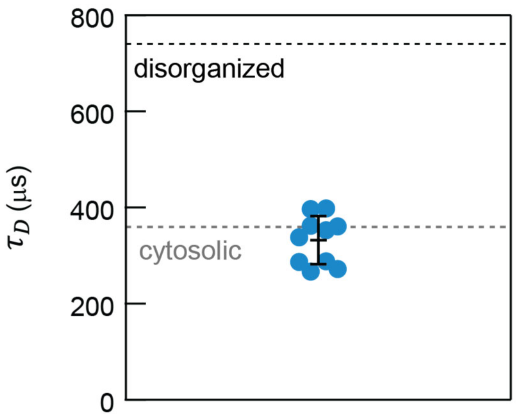


**Figure S3 Measurement error in cytoplasmic extracts**
Estimation of the upper bound of measurement error in cytoplasmic extracts. The data show 10 repetitive measurements at the same position in a mature border; each measurement was acquired identically as Fig. 2C and E. The error bar shows the standard deviation (50 μs). Note that since cytoplasmic extracts are highly dynamic and non-stationary (unlike homogeneous samples), this error includes measurement error and fluctuations of cytoplasmic organization.

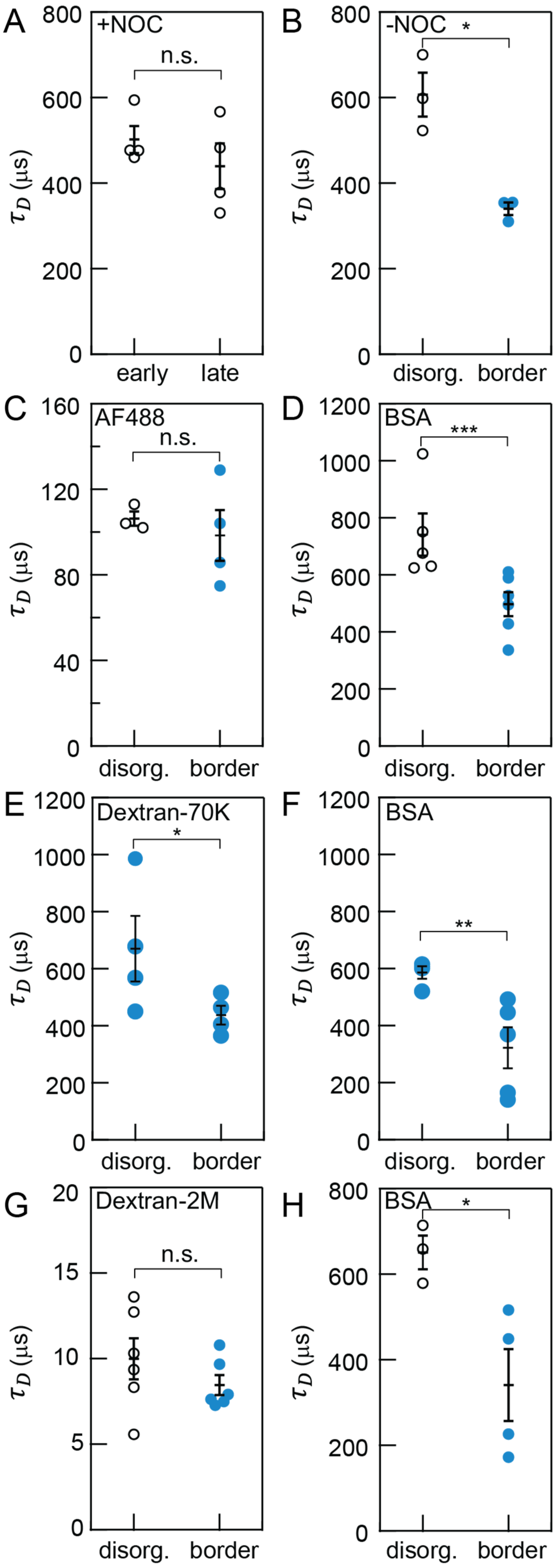


**Figure S4 Statistical tests**

Mann–Whitney U tests for data in Fig. 3C and 4. Asterisks (*) denotes the significance levels calculated as two-tailed p-values: ***, $p \leq 0.01$; **, $p \leq 0.05$; *, $p \leq 0.15$; n.s., not significant, $p > 0.15$. The exact p-values for (A-H) are 0.686, 0.100, 0.686, 0.004, 0.114, 0.016, 0.310, and 0.057, respectively. Error bars, S.E.M.

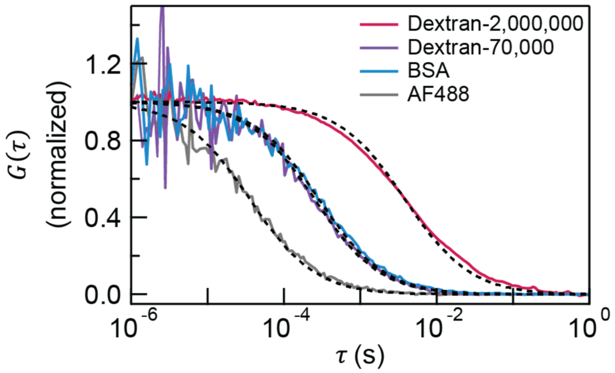


**Figure S5 Size estimation of probes used in this study**

The relative sizes of the diffusion probes were measured in PBS buffers using FCS. The autocorrelation functions were fitted by a Brownian model ($\alpha = 1$) (dashed curves). The diffusion times yield the relative Stokes radii by the Stokes-Einstein equation, $D \propto 1/R$. The relative Stokes radii for AF488, BSA, dextran-70K, and dextran-2M are 0.12, 1.00, 0.88, and 13.3, respectively. The relative values were converted to absolute lengths by taking the Stokes radius of BSA to be 3.5 nm. As a note, the polydisperse nature of dextran-2M could also be effectively fitted by the anomalous diffusion model, though the diffusion was expected to be Brownian (*6*). Nevertheless, the fitted diffusion times between the two models were similar.
